# Decoding of cortex-wide brain activity from local recordings of neural potentials

**DOI:** 10.1101/2021.10.14.464468

**Authors:** Xin Liu, Chi Ren, Zhisheng Huang, Madison Wilson, Jeong-Hoon Kim, Yichen Lu, Mehrdad Ramezani, Takaki Komiyama, Duygu Kuzum

## Abstract

**Objective:** Electrical recordings of neural activity from brain surface have been widely employed in basic neuroscience research and clinical practice for investigations of neural circuit functions, brain-computer interfaces, and treatments for neurological disorders. Traditionally, these surface potentials have been believed to mainly reflect local neural activity. It is not known how informative the locally recorded surface potentials are for the neural activities across multiple cortical regions.

**Approach:** To investigate that, we perform simultaneous local electrical recording and wide-field calcium imaging in awake head-fixed mice. Using a recurrent neural network model, we try to decode the calcium fluorescence activity of multiple cortical regions from local electrical recordings.

**Main results:** The mean activity of different cortical regions could be decoded from locally recorded surface potentials. Also, each frequency band of surface potentials differentially encodes activities from multiple cortical regions so that including all the frequency bands in the decoding model gives the highest decoding performance. Despite the close spacing between recording channels, surface potentials from different channels provide complementary information about the large-scale cortical activity and the decoding performance continues to improve as more channels are included. Finally, we demonstrate the successful decoding of whole dorsal cortex activity at pixel-level using locally recorded surface potentials.

**Significance:** These results show that the locally recorded surface potentials indeed contain rich information of the large-scale neural activities, which could be further demixed to recover the neural activity across individual cortical regions. In the future, our cross-modality inference approach could be adapted to virtually reconstruct cortex-wide brain activity, greatly expanding the spatial reach of surface electrical recordings without increasing invasiveness. Furthermore, it could be used to facilitate imaging neural activity across the whole cortex in freely moving animals, without requirement of head-fixed microscopy configurations.

## Introduction

As an important tool for electrophysiological recordings, neural electrodes implanted on the brain surface have been instrumental in basic neuroscience research to study large-scale neural dynamics(1) in various cognitive processes, such as sensorimotor processing(2) as well as learning and memory(3). In clinical settings, neural recordings have been adopted as a standard tool to monitor the brain activity in epilepsy patients before surgery for detection and localization of epileptogenic zones initiating seizures(4) and functional cortical mapping(5). Neural activity recorded from the brain surface exhibits rich information content about the collective neural activities reflecting the cognitive states and brain functions, which was leveraged for various types of brain-computer interfaces during the past decade. For example, surface potential recordings have been used for studying motor control, such as controlling a screen cursor(6) or a prosthetic hand(7). They have also been used to decode the mood of epilepsy patients, paving the way for the future treatment of neuropsychiatric disorders(8). Recent advances have shown that electrical recordings from cortical surface combined with the recurrent neural networks can even enable speech synthesis(9), demonstrating superior performance compared to those achieved through traditional noninvasive methods.

For the interpretation of surface potentials in terms of their neural correlates, most research has focused on local neural activities. The high-gamma band has been found to correlate with the ionic currents induced by synchronous synaptic input to the underlying neuron population(10). Besides that, the dendritic calcium spikes in the superficial cortical layers also contribute to surface potentials(11). Recently, it has been reported that even the action potentials of superficial cortical neurons could be detected in surface recordings(12). Despite the predominant focus of relating the surface potentials to local neural activity, they may also correlate with the large-scale activity of multiple cortical regions. This could be achieved through the intrinsic correlations of the spontaneous activities among large-scale cortical networks(13, 14) due to the anatomical connectivity(15) and the global modulation of neuromodulatory projections(16). However, this rich information content of surface potentials encoded for the large-scale cortical activity remains unexploited and little is known about how local surface potentials are correlated with the spontaneous neural activities of distributed large-scale cortical networks.

In this work, we investigate whether the rich information content of the local neural potentials recorded from brain surface can be harnessed to infer the cortex-wide brain activity. We employed optically transparent graphene microelectrodes implanted over the mouse somatosensory cortex and posterior parietal cortex to perform simultaneous wide-field calcium imaging of the entire dorsal cortex during local neural recordings in awake mice. Multimodal datasets generated by these experiments were used to train a recurrent neural network model to learn the hidden spatiotemporal mapping between the local surface potentials and the cortex-wide brain activity detected by wide-field calcium imaging. We demonstrated that both the average spontaneous activity from multiple cortical regions and the pixel-level cortex-wide brain activity can be inferred from locally recorded surface potentials. Our results show that in addition to the changes of local neural activity, the spontaneous fluctuations of locally recorded surface potentials also reflect the collective variations of large-scale neural activities across the entire cortex.

## Methods

### Fabrication of graphene array

Electrode arrays were fabricated on 4” Silicon wafers spin coated with 20 μm-thick PDMS. 50 μm-thick PET (Mylar 48-02F-OC) was placed on the adhesive PDMS layer and used as the array substrate. 10 nm of chromium and 100 nm of gold were deposited onto the PET using a Denton 18 Sputtering System. The metal wires were patterned using photolithography and wet etching methods. Single-layer graphene was placed on the array area using a previously developed transfer process(17, 18). The wafer was then soft baked for 5 minutes at 125°C to better adhere graphene to the substrate. PMMA was removed via a 20-minute acetone bath at room temperature then rinsed with isopropyl alcohol and DI water for ten 1-minute cycles. The graphene channels were patterned using AZ1512/PMGI bilayer photolithography then oxygen plasma etched (Plasma Etch PE100). A four-step cleaning method was performed on the array consisting of an AZ NMP soak, remover PG soak, acetone soak, and 10-cycle isopropyl alcohol/DI water rinse. 8 μm-thick SU-8 2005 was spun onto the wafer as an encapsulation layer and openings were created at the active electrical regions using photolithography. The array was then given a final 10-cycle isopropyl alcohol/DI water rinse to clean SU-8 residue and baked for twenty minutes at temperature progressing from 125°C to 135°C.

### Animals

All procedures were performed in accordance with protocols approved by the UCSD Institutional Animal Care and Use Committee and guidelines of the National Institute of Health. Mice (cross between CaMKIIa-tTA:B6;CBA-Tg(Camk2a-tTA)1Mmay/J [JAX 003010] and tetO-GCaMP6s: B6;DBA-Tg(tetO-GCaMP6s)2Niell/J [JAX 024742], Jackson laboratories) were group-housed in disposable plastic cages with standard bedding in a room with a reversed light cycle (12 h-12 h). Experiments were performed during the dark period. Both male and female healthy adult mice were used. Mice had no prior history of experimental procedures that could affect the results.

### Surgery and multimodal experiments

Adult mice (6 weeks or older) were anesthetized with 1–2% isoflurane and injected with baytril (10 mg/kg) and buprenorphine (0.1 mg/kg) subcutaneously. A circular piece of scalp was removed to expose the skull. After cleaning the underlying bone using a surgical blade, a custom-built head-bar was implanted onto the exposed skull over the cerebellum (∼1 mm posterior to lambda) with cyanoacrylate glue and cemented with dental acrylic (Lang Dental). Two stainless-steel wires (791900, A-M Systems) were implanted into the cerebellum as ground/reference. A craniotomy (∼7 mm × 8 mm) was made to remove most of the dorsal skull and the graphene array was placed on the surface of one hemisphere, covering somatosensory cortex (S1) and posterior parietal cortex (PPC). The exposed cortex and the array were covered with a custom-made curved glass window, which was further secured with Vetbond (3M), cyanoacrylate glue and dental acrylic. Animals were fully awake before recordings. During recording, animals were head-fixed under the microscope, free to run or move their body, and not engaged in task.

The wide-field calcium imaging was performed using a commercial fluorescence microscope (Axio Zoom.V16, Zeiss, objective lens (1x, 0.25 NA)) and a CMOS camera (ORCA-Flash4.0 V2, Hamamatsu) through the curved glass window as previously described(19). The light source for wide-field calcium imaging is HXP 200 C (Zeiss). The filter set for imaging GCaMP signals is commercially installed in the microscope. It consists of a bandpass filter for the excitation light (485 ± 17 nm), a beamsplitter (500 nm), and a tunable bandpass filter centered at 520 nm for the emission light. Images were acquired using HCImage Live (Hamamatsu) at 29.98 Hz, 512 × 512 pixels (field of view: 8.5 mm x 8.5 mm, binning: 4, 16 bit).

The microelectrode array was connected to a custom-made connector board through a ZIF connector. The surface potential data was sampled with Intan RHD2132 amplifier and recorded using Intan RHD2000 system. The sampling frequency was 10 kHz. To synchronize the electrical recording with the calcium imaging, we used a trigger signal (TTL), a 2 V pulse of 1 s, to trigger the start of the calcium imaging. Meanwhile, this trigger signal was also sent to the ADC of Intan recording system. During the data processing stage, we detected the onset of the pulse and aligned the imaging data and electrical data to that time point. Three mice were recorded, each having 2-3 recording sessions. The length for each recording session was 1 hour.

### ΔF/F processing

To obtain the ΔF/F time series from the wide-field calcium imaging data, we first down-sampled the 512 × 512 pixel images to smaller images of 128 × 128 pixels. For each pixel, we defined a dynamic fluorescence (F) baseline for a given time point as the 10th percentile value over 180 s around it. For the beginning and ending of each imaging block, the following and preceding 90-s window was used to determine the baseline, respectively. An 8th order 6 Hz Butterworth low-pass filter was applied to the ΔF/F activity of each pixel to remove the high frequency noise and hemodynamic contamination from heartbeat. The activity of each cortical region was obtained by averaging over the ΔF/F signals from all the pixels within the same cortical regions defined by the Allen Brain Atlas.

### Surface recording data processing

The raw surface recording data was first passed through notch filters to eliminate the 60 Hz powerline contaminations and their higher harmonics at 120 Hz and 180 Hz. The signals were further filtered with multiple 6th order Butterworth band-pass filters designed for different frequency bands (δ: 1 – 4 Hz, θ: 4 – 7 Hz, α: 8 – 15 Hz, β : 15 – 30 Hz, γ : 31-59 Hz, H-γ : 61 – 200 Hz). The resulting signals were squared and smoothed by a Gaussian function with 100 ms time window to obtain an estimate of the instantaneous power. To prepare the input data for the decoding neural network, the power traces at different frequency bands were down-sampled to 29.98 Hz by interpolation to match the sampling rate of calcium imaging data. To suppress the potential artifacts in the recording signal, at each frequency band we clip the power traces with a threshold of 95 percentile.

### Neural network models

The neural network model consists of a sequential stacking of a linear hidden layer, one bidirectional LSTM layer and a linear readout layer. The first linear layer was followed by batch normalization, ReLU activation, and dropout with a probability of 0.3. The LSTM layer was followed by batch normalization. The multichannel power at different frequency bands were used as inputs to the network. To decode the neural activity at each time step t, the power segments between [t-1.5s, t+1.5s] was used (90 time steps in total). The first linear layer had 16 neurons and the bidirectional LSTM had 8 hidden neurons. The same neural network model was used for the two decoding tasks except that the number of neurons in the final output layer differs based on the targeting output. To decode the ΔF/F activity of 12 cortical regions simultaneously, the output neuron number was set to 12. To decode the cortex-wide brain activity, the output neuron number was set to 10 to generate the scores for the 10 ICs. Assuming using 6 frequency bands from 16 recording channels, setting sequence length of LSTM layer to 90, and setting batch size to 128, the input and output size for each layer of the model is shown in Table 1. Note that we flattened the last two dimensions of the LSTM output to make it 128 × 1440 before feeding it to the last linear layer.

**Table 1.** The size for input and output tensors of each layer.

|  | Input size | Output size |
| --- | --- | --- |
| First linear layer | 128 x 90 x 96 | 128 x 90 x 16 |
| Bi-LSTM layer | 128 x 90 x 16 | 128 x 90 x 16 |
| Last linear layer | 128 x 1440 | 128 x 12 or 128 x 10 |

The neural network model was implemented in Pytorch(20). The model parameters were trained through Adam optimizer with learning rate = 1e-4, beta1 = 0.9, beta2 = 0.999, epsilon = 1e-8. The batch size was 128 and the training usually converged within ∼30 epochs. For both tasks, the mean squared error was chosen as the loss function. We performed 10-fold cross-validation where each 1 h recording session was chunked into ten segments, each lasting for 6 min. The neural network model was trained on 9/10 of the data segments and tested on a different held-out segment that was unseen during the training. To evaluate the model performance, correlation between the decoded and ground truth data for each held-out set was averaged. For each 1 h recording session, a new network model is trained and tested. Then, for each mouse, the correlation was further averaged across the recording sessions to give the performance for that mouse.

### Statistical tests

All statistical analyses were performed in MATLAB. Statistical tests were two-tailed and significance was defined by alpha pre-set to 0.05. All the statistical tests are described in the figure legends. Multiple comparisons were corrected for by Benjamini-Hochber corrections.

## Results

### Multimodal recordings of cortical activity

Cortical recordings in both clinical applications and neuroscience studies use conventional metal-based neural electrode arrays. However, these opaque neural electrodes are not suitable for multimodal recordings combined with optical imaging since they will block the field of view and generate light-induced artifacts under optical imaging(21, 22). Compared to conventional neural electrode arrays, graphene-based surface arrays are optically transparent and free from light-induced artifacts, both of which are key to the simultaneous electrical recordings and optical imaging of cortical activity(18, 23). Wide-field calcium imaging is an optical imaging technique that can provide simultaneous monitoring of large-scale cortical activity and has been used to study the dynamics of multiple cortical regions and their coordination during behavior(19, 24-26). Compared to fMRI that also offers large spatial coverage, the wide-field calcium imaging provides better spatiotemporal resolution and higher signal-to-noise ratio(25). It has been shown that wide-field calcium signals mainly reflect local neural activity(19). Therefore, the multimodal experiments combining electrical recordings based on graphene arrays and the wide-field calcium imaging generate unique datasets that are ideal for investigating the mapping from local neural signals to large-scale cortical activity.

We fabricated transparent graphene arrays on 50 μm thick flexible polyethylene terephthalate (PET) substrates(18, 23) (see methods for details). 10 nm of chromium and 100 nm of gold were deposited onto the PET and the metal wires were patterned using photolithography and wet etching methods. The graphene layer was transferred and patterned with photolithography and oxygen plasma etching to form electrode contacts. Finally, 8 μm-thick SU-8 was used as an encapsulation layer and openings were created at the active electrical regions using photolithography. The graphene array has 16 recording channels, each of size 100 × 100 μm. The spacing between adjacent channels is 500 μm. The graphene array was implanted unilaterally over the somatosensory cortex (S1) and posterior parietal cortex (PPC) of the mice to perform the simultaneous electrical recordings and wide-field calcium imaging (Figure 1a). We performed multimodal recordings of spontaneous neural activity in awake mice during either quiet resting state or actively running or moving. An example wide-field image obtained during the experiment is shown in Figure 1b. Note that the cortical activity under the array could still be observed due to the transparency of the graphene electrode. Based on Allen brain atlas, we parcellated the brain into 12 different ipsilateral (the hemisphere with array implanted) and contralateral cortical regions (Figure 1c), including the primary and secondary motor cortices (M1, M2), the somatosensory cortex (S1), the posterior parietal cortex (PPC), the retrosplenial cortex (RSC), and the visual cortex (Vis). Representative spontaneous cortical activity recorded during the experiment is shown in Figure 1d. We observed dynamical changes of large-scale cortical activity, involving co-activations of multiple cortical regions. In the simultaneous multi-channel neural recordings, we also observed differences in power traces from different channels at multiple frequency bands during the spontaneous cortical activity (Figure 1e). Compared with the fluorescence activity, the neural potential signal has a much higher temporal resolution and richer frequency components.

**Figure 1.**
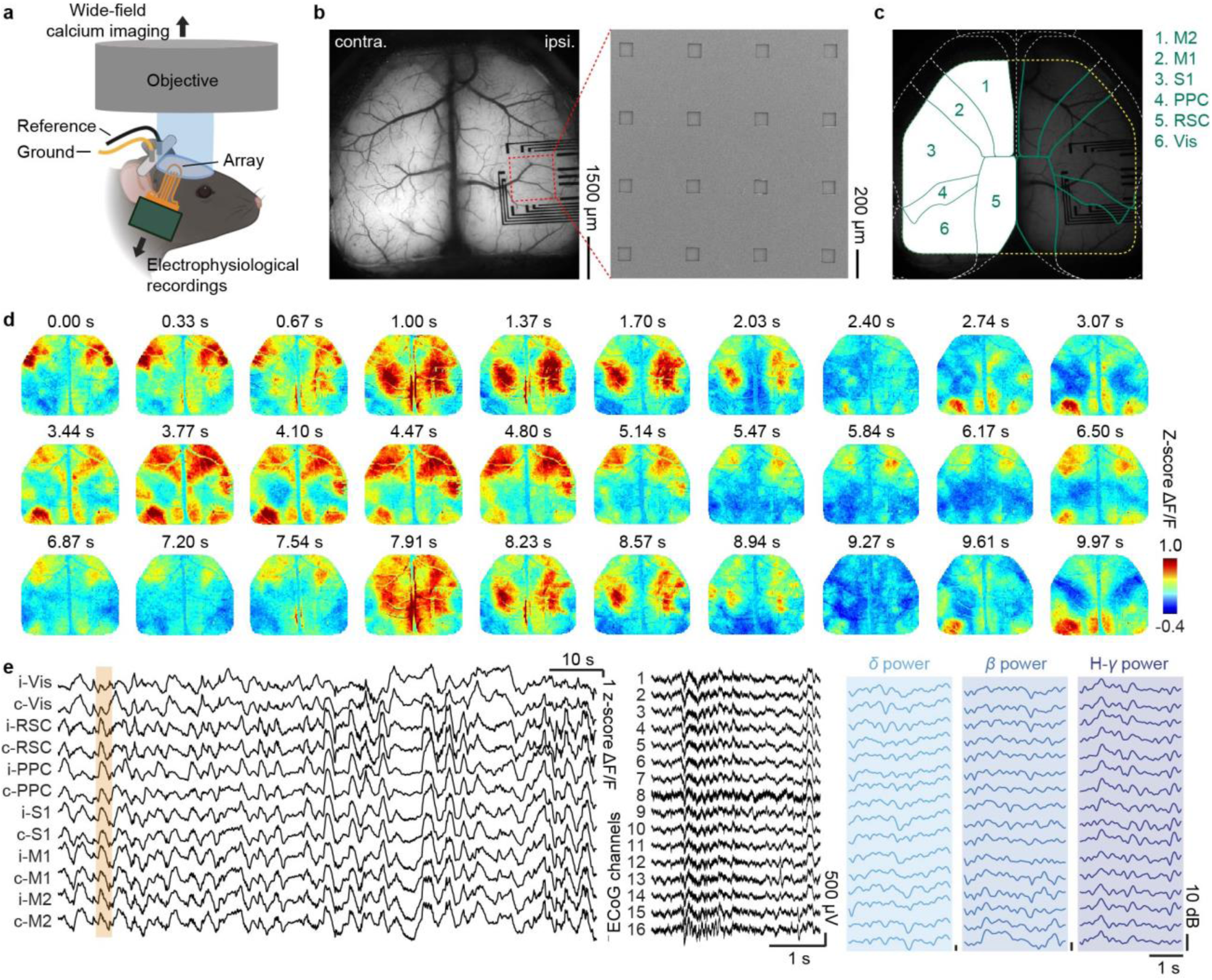
Simultaneous multimodal wide-field calcium imaging and surface potential recordings. a, Schematic of the multimodal experimental setup combining neural recordings using transparent graphene electrodes and wide-field calcium imaging. b, Example field of view of wide-field calcium imaging during experiment (left). Clear area at the center of the transparent array includes 16 graphene electrodes, whose scanning electron microscope image is shown on the right. c, Imaged cortical regions based on Allen Brain Atlas. M2: secondary motor cortex; M1: primary motor cortex; S1: primary somatosensory cortex; PPC: posterior parietal cortex; RSC: retrosplenial cortex; Vis: visual cortex. d, Wide-field fluorescence activity during 10-s long recordings, showing the diverse spontaneous activity across the mouse cortex. e, Fluorescence activity for different cortical regions (left), the simultaneously recorded neural signals (middle) for a 3-s time interval (marked by the yellow bar on the left), and their power at three frequency bands (δ: 1 - 4 Hz, β : 15 – 30 Hz, H-γ : 61 – 200 Hz, right three columns).

### Cortical activity decoder design

To investigate whether the locally recorded surface potentials could be used to infer the cortex-wide brain activity, we investigated two decoding tasks, namely the decoding of the average activity from individual cortical regions and the decoding of pixel-level cortex-wide brain activity. To achieve these goals, we developed a compact neural network model consisting of a linear hidden layer, a one-layer LSTM network, and a linear readout layer (Figure 2, See methods for details). In both tasks, the signal power traces of multiple frequency bands recorded from different recording channels were used as inputs to the neural network. In the first task, the neurons in the output layer of the neural network directly generate the activity of all the cortical regions simultaneously. In the second task, we first performed principal component analysis (PCA) on the cortical activity to remove the noise and reduce the dimensionality of the data. Across all the mice, the top 10 principal components (PCs) explain > 92% variance in the data (Supplementary Figure 1). Then based on the PCA results, we further performed spatial independent component analysis (ICA) to obtain the independent components (ICs) and their weighting scores for the data at each time frame. In all the three mice, the identified ICs reflect different functional modules and hemodynamic signals on blood vessels (Supplementary Figure 2) and provide a set of functionally meaningful basis for the decomposition of the large-scale cortical activity. The output layer of the neural network directly generates the estimated weighting scores of individual ICs, which were further used to reconstruct the cortex-wide brain activity at each time frame with pixel-level spatial resolution (Figure 2).

**Figure 2.**
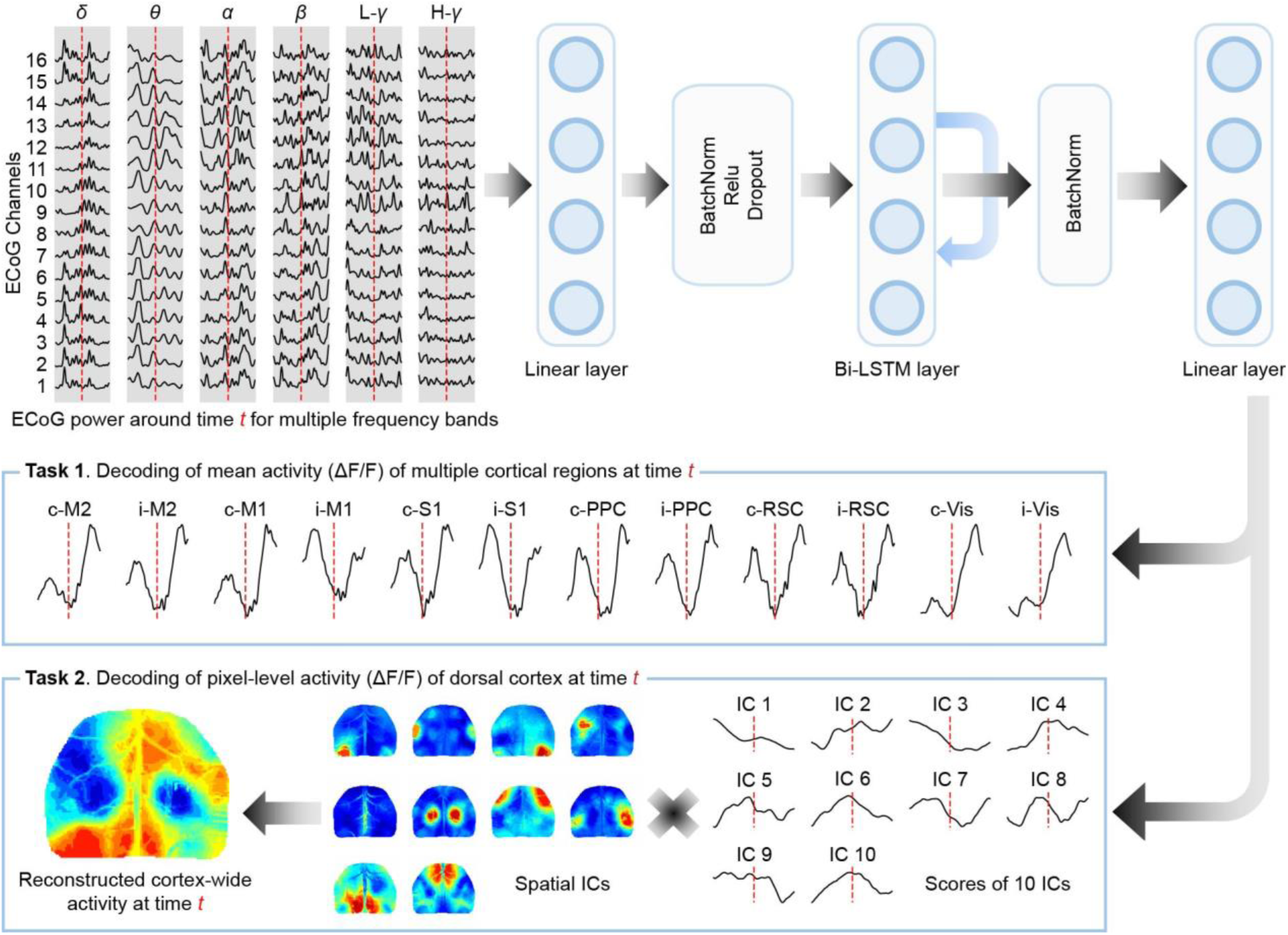
Schematic of the decoding model. Signal power from different channels during time interval [t-1.5 s, t+1.5 s] (90 time steps) is used to decode the cortical activity at time point t. The decoding neural network model consists of a sequential stacking of a linear hidden layer, one bidirectional LSTM (Bi-LSTM) layer and a linear readout layer. For the task of decoding the mean ΔF/F activity from multiple cortical regions, the final linear readout layer directly outputs the activities of 12 cortical regions at time t. For the task of decoding the pixel-level cortex-wide brain activity, the final linear readout layer outputs the weighting scores for all the independent components at time t, from which the cortex-wide brain activity at time t is reconstructed.

### Decoding of activity for individual cortical regions

Based on the multimodal data we collected during the animal experiment and the above designed decoder network model, we decoded the mean activity of both the ipsilateral and contralateral cortical regions using the power of six frequency bands from all recording channels. An example of decoded and ground truth (ΔF/F from wide-field calcium imaging) cortical activity from one held-out set is shown in Figure 3a. The decoding performances for S1, PPC, and RSC regions closely resemble the ground truth cortical activity, while the decoding performances for M1, M2, and Vis are lower, possibly due to their increasing distances to the recording electrode array. We performed the same decoding analysis using shuffled data. The results show decoding performance close to zero (Supplementary Figure 3a). We evaluated the stability of the decoding performance across time using a 30 s sliding window. The results show that the decoding performance fluctuates from time to time but remains stable in the longer time intervals (Supplementary Figure 4A). We also compared the decoding performance of individual cortical regions during rest and movement intervals and found similar decoding performance between rest and movement phases (Supplementary Figure 4B). Therefore, the fluctuations of the decoding performance across time are not due to animal movements.

**Figure 3.**
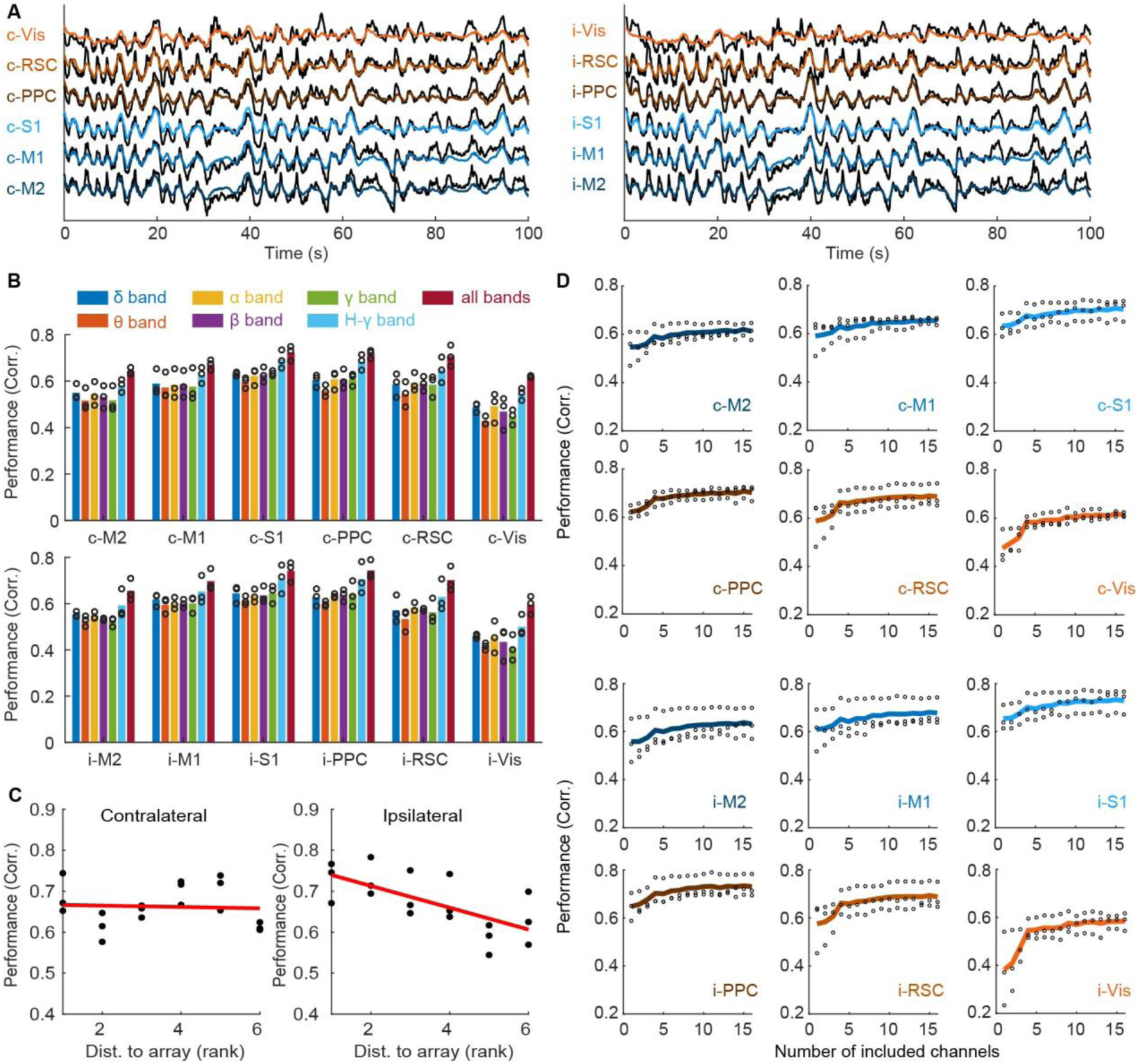
Decoding the activities of multiple cortical regions. a, Decoded (colorful) v.s. ground truth (black) ΔF/F activity of different cortical regions in the contralateral (left) and ipsilateral (right) hemispheres for one mouse. b, Decoding performance evaluated for different cortical regions in the contralateral (top) and ipsilateral (bottom) hemispheres using different frequency bands (δ: 1 – 4 Hz, θ: 4 – 7 Hz, α: 8 – 15 Hz, β : 15 – 30 Hz, γ : 31-59 Hz, H-γ : 61 – 200 Hz, and all 6 frequency bands). Each dot marks the mean correlation evaluated by 10-fold cross-validation using the data recorded from one mouse. c, Decoding performance for different cortical regions in the contralateral (left) and ipsilateral hemispheres evaluated as a function of distance (rank orders). Each dot is the mean correlation for one mouse given by 10-fold cross-validation. For ipsilateral hemisphere, the decoding performance decreases as the distance rank to the electrode array increases (ρ = -0.676, P = 0.002, n = 18). For contralateral hemisphere, no such correlation is observed (ρ = -0.163, P = 0.519, n = 18). Distances from the center of the array to the center of each cortical region: i-M2 3.63 mm, i-M1 2.65 mm, i-S1 0.98 mm, i-PPC 0.7 mm, i-RSC 2.36 mm, i-Vis 2.49 mm, c-M2 5.01 mm, c-M1 5.53 mm, c-S1 5.96 mm, c-PPC 5.37 mm, c-RSC 3.83 mm, c-Vis 6.32 mm. d, Decoding performance for different cortical regions in the contralateral (top) and ipsilateral (bottom) hemispheres using all the frequency bands, but different numbers of recording channels. Each dot marks the mean 10-fold cross-validated correlation over all the recording sessions for one mouse. Each line is the mean correlation averaged across 3 mice. For all the cortical regions, the decoding performance increases as more recording channels are included (P<0.05, n = 48, FDR correction).

To further evaluate how informative different frequency bands are for the decoding of the activity from different cortical regions, we used the signal power from different frequency bands of all channels as inputs and performed 10-fold cross-validation to evaluate the decoding performance of the neural network model. We find that even though all the frequency bands are informative of the activities in different cortical regions, the high gamma power band gives the highest decoding performance for all the cortical regions compared to other frequency bands (Supplementary Figure 5 and 6). However, across all the cortical regions, using all the frequency bands yields the best decoding performance compared to using any single frequency band (Figure 3b), implying that different frequency bands provide complementary information about the activity in multiple cortical regions. Decoding with the shuffled data gives performance close to zero for all the frequency bands (Supplementary Figure 3b). For the ipsilateral cortical regions, we also find a negative correlation between their decoding performance and their distance ranks to the recording array. However, for the contralateral cortical regions, no significant correlation is observed (Figure 3c). When comparing the decoding results of the activity from ipsilateral cortical regions using different frequency bands, we find that higher frequency bands tend to have a steeper slope for the decoding performance vs. distance to the recording array (Supplementary Figure 7).

Besides the frequency bands, we also examined whether different recording channels encode nonredundant information for decoding the activity of different cortical regions. Therefore, we evaluated the decoding performance of the neural network model using all six frequency bands from different numbers of channels. Specifically, we performed 10-fold cross-validation on the neural network multiple times and each time we sequentially added the signal power of all frequency bands from one random channel until all the channels were included. As shown in Figure 3d, for all the cortical regions, increasing the number of channels significantly improves the decoding performance, suggesting that recording channels of local neural potentials provide nonredundant information about the activity from multiple cortical regions. On the other hand, decoding with the shuffled data gives performance close to zero for different number of included channels (Supplementary Figure 3c).

### Decoding of pixel-wise activity across cortex

Given that the local neural signals encode average activity from individual cortical regions, which could be recovered by the neural network model using multi-channel signal power of different frequency bands, we further investigated whether the pixel-level activity across the whole dorsal cortex could also be decoded using locally recorded neural signals. The same neural network model for decoding the average activity in different cortical regions was then employed to simultaneously decode the ten IC scores at each time frame. The power traces of all the six frequency bands from all the recording channels were used as inputs to the neural network. An example of the decoded and ground truth scores for the ten ICs from one held-out set is shown in Figure 4b. The decoding result using shuffled data is shown in Supplementary Figure 3d. Based on the decoded IC scores and the IC modules (Figure 4a), the pixel-level cortex-wide activity at each time frame could be reconstructed. Examples of the reconstructed pixel-level cortex-wide activity during 4 representative time intervals are shown in Figure 4c. The reconstructed cortex-wide activity captured various patterns of cortical activations in ground truth, including both the synchronous and asynchronous activations among different cortical regions. These diverse activation patterns cannot be explained solely by PC1 (see Figure 4c and the supplementary videos). To further quantify this observation, we computed the correlation between the ground truth activity of each ICs and the PC1. The median correlations between IC1, IC2 and IC8 to PC1 are close to zero, showing that their activities are not strongly correlated to PC1 (Supplementary Figure 8). These results suggest that the model does not merely predict dominant activity patterns showing activation around S1 and RSC. We found that all the ten IC scores could be decoded using the locally recorded neural signals (Figure 4d, Supplementary Figure 9). We demonstrated that the pixel-level cortex-wide activity could be reconstructed for all the recording sessions (Supplementary Videos 1-7). This reveals that the cortical activations of distinct functional modules indeed induce different responses in local cortical electrical signals, which could be in turn used to recover the diverse cortex-wide activity patterns. In addition to cortical activity, in all the mice, we observed one or two ICs showing the hemodynamic activity (Supplementary Figure 2). Our decoding results also show that these hemodynamic activities could be decoded from the neural recordings, which is mainly due to the fact that hemodynamic activity and the neural activity are often correlated(27). Next, we examined the pixel-level correlations between the decoded and ground truth activities imaged using wide-field imaging in individual cortical regions. We observed high correlations between the decoded and the ground truth data for all cortical regions (Figure 4e) and close-to-zero correlations using shuffled data (Supplementary Figure 3e). The activities of cortical regions closer to the array are better decoded than those of the cortical regions far away from the array. Consistent with the decoding of mean activity in each cortical region, the pixel-wise correlation decreases as the distance rank to the surface array increases for the ipsilateral cortical regions, whereas for the contralateral cortical regions no such correlation exists (Figure 4f).

**Figure 4.**
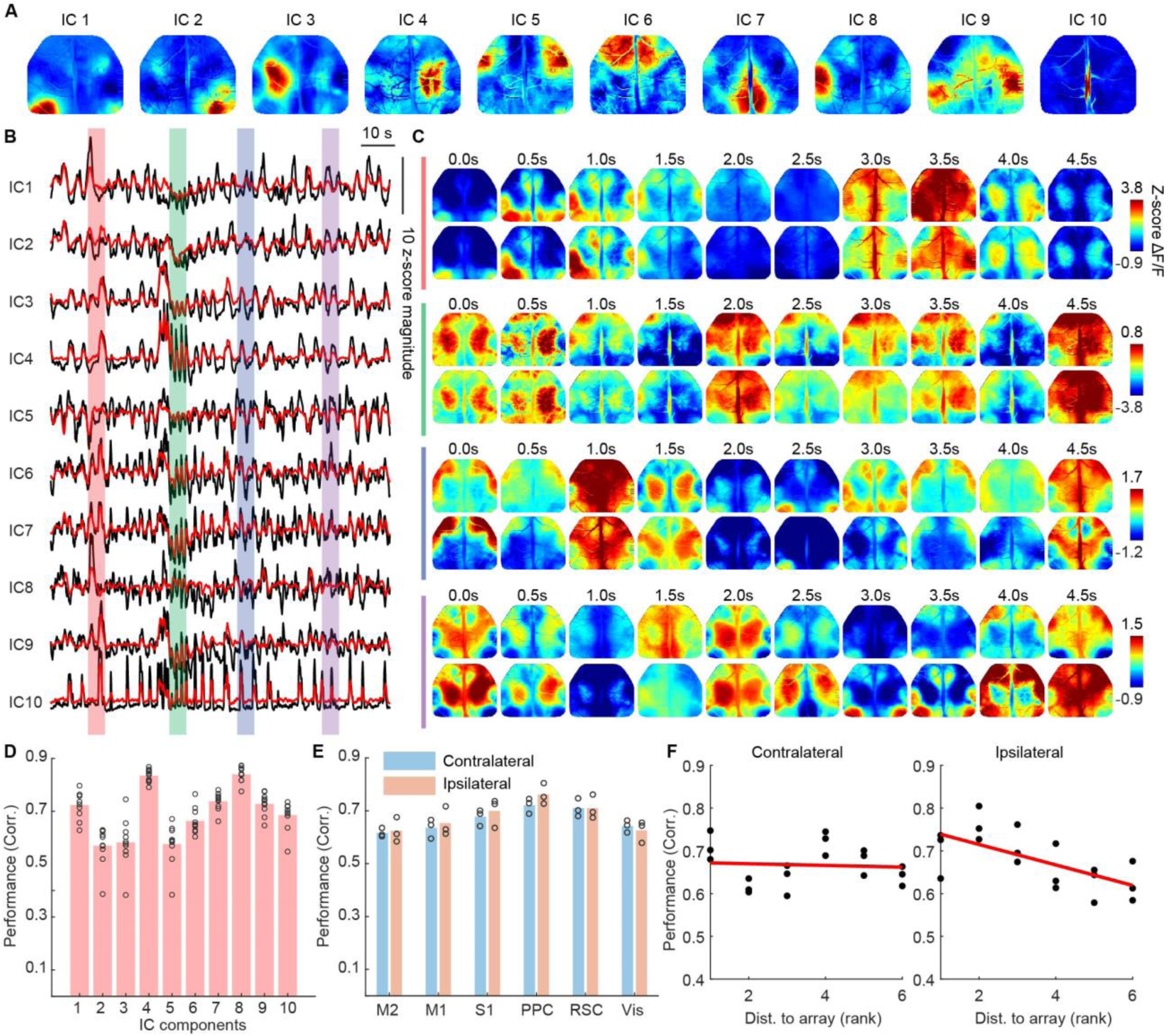
Decoding of the pixel-level cortex-wide brain activity. a, Identified independent components for the cortical activities recorded in one mouse, showing different functional modules of the cortical activity (IC 1-9) and the blood vessel activity (IC 10). b, Decoded (red) and ground truth (black) weighting scores of the observed cortex-wide activity onto the 10 ICs shown in a. c, Reconstructed (top rows) and ground truth (bottom rows) cortex-wide ΔF/F activity for 4 different time intervals, each lasting for 5 s, as indicated with different colors in b. For visualization, the reconstructed and true cortex-wide brain activity are shown for every 0.5 s. d, Decoding performance evaluated for different ICs for one recording session. Each dot marks the decoding performance evaluated on one fold during the 10-fold cross-validation. The weighting scores for all the 10 ICs could be successfully decoded. e, Decoding performance evaluated at pixel-level for all the cortical regions in the ipsilateral and contralateral hemispheres. Each dot marks the mean 10-fold cross-validated correlation for individual pixels of one specific cortical region from one mouse. f, Pixel-wise decoding performance evaluated at individual cortical regions and displayed as a function of distance to the array (rank orders). For ipsilateral hemisphere, the decoding performance decreases as the distance to the electrode array increases (ρ = -0.649, P = 0.003, n = 18). For contralateral hemisphere, no correlation is observed (ρ = -0.074, P = 0.770, n = 18).

## Discussion

In this work, we performed multimodal recordings of local neural potentials and wide-field calcium imaging in awake mice and developed a recurrent neural network model to decode the large-scale spontaneous cortical activity from the locally recorded multi-channel electrical signals. Both the averaged and the pixel-level activity across the entire dorsal cortex could be decoded, and the best decoding performance was achieved using all frequency bands of recorded neural potentials. These results suggest that even though the cortical electrical recording is a complex signal contributed by various mechanisms at multiple spatial scales, the responses in individual frequency bands across multiple recording channels still provide important discriminative information about the activity of different cortical regions. By developing a decoder model, the mixed information in the electrical signal responses could be used to recover the simultaneously recorded cortex-wide brain activity.

The cortical potentials have long been believed to mainly detect local neural activities that are within a sensing distance between 500 μm to 1-3mm(28-30), depending on the size of the electrode(28) as well as the spatial correlation pattern of neural activity(29). Consistent with this claim, for the decoding of mean activity from individual cortical regions, we find a decreasing decoding performance for the ipsilateral cortical regions located ∼1.5-3mm from the array. Interestingly, for the contralateral cortical regions, the decoding is still possible even though their activities are unlikely to be directly detected by the neural electrodes. We suspect that the successful decoding of contralateral cortical regions is mainly due to the fact that the spontaneous activities of same functional cortical regions in both hemispheres are often correlated (Supplementary Figure 10). Such correlated activity could arise from the anatomical connectivity(15) and further orchestrated by neuromodulatory projections(16).

Our decoding results for the activity of individual cortical regions show that even with single recording channel, the decoding is possible (mean correlation performance between 0.35-0.65 for different cortical regions). By including more channels, initially we observed an increase in decoding performance, but the performance starts to saturate after the inclusion of ∼10 recording channels (mean correlation performance between 0.6-0.75 for different cortical regions). We suspect that this is mainly because of the fact that the neural potentials in adjacent channels are partially correlated due to the volume conduction in the brain tissue(31, 32). It has been shown that the correlation between neural potentials from adjacent channels at different frequency bands decreases as the distance increases(33, 34). Even though the cross-channel correlation at high frequency bands is lower than that at low frequency bands, it does not go below chance level even with a distance of ∼1.5mm. However, our results empirically confirm that even though the neural potentials from adjacent channels are partially correlated, they still differentially encode information about the cortical activities to some extent so that sequentially including more recording channels tends to increase the decoding performance. However, beyond a certain threshold adding more channels does not further increase the decoding performance.

For the decoding of cortex-wide brain activity, instead of attempting to directly reconstruct the activity of individual pixels, we chose to perform PCA followed by spatial ICA on the cortical activity and later to decode IC scores to recover the cortex-wide activity at pixel level. The adoption of this approach was based on both scientific and computational considerations. First, the PCA effectively reduced the spatial dimensions, while preserving a large proportion of variance in cortical activity. Since the activity of each single pixel was noisy, performing PCA reduced the noise, leading to a more reliable estimate of the true activity. Second, choosing the IC scores as network outputs greatly reduced the parameters in the output layer of the neural network model, prevented overfitting, and speeded up model training. Finally, the spontaneous cortex-wide brain activity was decomposed into a set of local and spatially organized cortical activation patterns based on neural activity, generating a biologically meaningful decomposition that matches the brain dynamics. This decomposition provides a good demixing of cortex-wide brain activity and enables a tractable mapping from cortical neural responses, which can be learned by the decoding network model. Taken together, these results reveal that the activation of different cortical functional modules are associated with distinct components in local neural activity. By exploiting the mapping between the two modalities, the decoding of cortex-wide brain activity is possible from locally recorded neural signals.

## Conclusion

In this paper, we designed a neural network model to show that both the mean activity of different cortical regions and the pixel-level cortex-wide neural activity can be decoded using locally recorded surface potentials. These findings demonstrated that the locally recorded neural potentials indeed contain rich information for large-scale neural activity and the surface potential responses in different frequency bands and different recording channels provide distinct information about the large-scale neural activity.

## Data and code availability

The data and custom Python and MATLAB codes are available on Github repository https://github.com/xinliuucsd/Cortex-wide-Fluorescence-Decoding.

## Acknowledgements

This research was supported by grants from the ONR Young Investigator Award (N00014161253), NSF (ECCS-2024776, ECCS-1752241 and ECCS-1734940) and NIH (R21 EY029466 and R21 EB026180 and DP2 EB030992) to DK, and grants from NIH (R01 NS091010A, R01 EY025349, R01 DC014690, R21 NS109722, and P30 EY022589), Pew Charitable Trusts, and David & Lucile Packard Foundation to T.K. Fabrication of the electrodes was performed at the San Diego Nanotechnology Infrastructure (SDNI) of UCSD, a member of the National Nanotechnology Coordinated Infrastructure, which is supported by the National Science Foundation (Grant ECCS-1542148).

## Author contributions

This work was conceived by XL, CR, and DK. Microelectrode array fabrication was performed by YL and MW. All animal experiments were performed by CR and XL and analyzed by XL and CR with contributions from TK and DK. Neural network implementation was done by XL and ZH. The manuscript was written by XL, CR, MW, JHK and DK and edited by all authors.

## Ethics declarations

The authors declare no competing interests.

## References

1. Muller L, Piantoni G, Koller D, Cash SS, Halgren E, Sejnowski TJ. Rotating waves during human sleep spindles organize global patterns of activity that repeat precisely through the night. Elife. 2016;5:e17267.

2. Cogan GB, Thesen T, Carlson C, Doyle W, Devinsky O, Pesaran B. Sensory–motor transformations for speech occur bilaterally. Nature. 2014;507(7490):94–8.

3. Khodagholy D, Gelinas JN, Buzsáki G. Learning-enhanced coupling between ripple oscillations in association cortices and hippocampus. Science. 2017;358(6361):369–72.

4. Osorio I, Frei MG, Wilkinson SB. Real-time automated detection and quantitative analysis of seizures and short-term prediction of clinical onset. Epilepsia. 1998;39(6):615–27.

5. Ojemann G, Ojemann J, Lettich E, Berger M. Cortical language localization in left, dominant hemisphere: an electrical stimulation mapping investigation in 117 patients. Journal of neurosurgery. 1989;71(3):316–26.

6. Schalk G, Miller KJ, Anderson NR, Wilson JA, Smyth MD, Ojemann JG, et al. Two-dimensional movement control using electrocorticographic signals in humans. J Neural Eng. 2008;5(1):75.

7. Chestek CA, Gilja V, Blabe CH, Foster BL, Shenoy KV, Parvizi J, et al. Hand posture classification using electrocorticography signals in the gamma band over human sensorimotor brain areas. J Neural Eng. 2013;10(2):026002.

8. Sani OG, Yang Y, Lee MB, Dawes HE, Chang EF, Shanechi MM. Mood variations decoded from multi-site intracranial human brain activity. Nature Biotechnology. 2018:954–61.

9. Anumanchipalli GK, Chartier J, Chang EF. Speech synthesis from neural decoding of spoken sentences. Nature. 2019;568(7753):493–8.

10. Ray S, Crone NE, Niebur E, Franaszczuk PJ, Hsiao SS. Neural correlates of high-gamma oscillations (60–200 Hz) in macaque local field potentials and their potential implications in electrocorticography. J Neurosci. 2008;28(45):11526–36.

11. Suzuki M, Larkum ME. Dendritic calcium spikes are clearly detectable at the cortical surface. Nature Communications. 2017;8:276.

12. Khodagholy D, Gelinas JN, Thesen T, Doyle W, Devinsky O, Malliaras GG, et al. NeuroGrid: recording action potentials from the surface of the brain. Nat Neurosci. 2015;18(2):310–5.

13. Nir Y, Mukamel R, Dinstein I, Privman E, Harel M, Fisch L, et al. Interhemispheric correlations of slow spontaneous neuronal fluctuations revealed in human sensory cortex. Nature neuroscience. 2008;11(9):1100–8.

14. Fox MD, Raichle ME. Spontaneous fluctuations in brain activity observed with functional magnetic resonance imaging. Nat Rev Neurosci. 2007;8(9):700–11.

15. O’Reilly JX, Croxson PL, Jbabdi S, Sallet J, Noonan MP, Mars RB, et al. Causal effect of disconnection lesions on interhemispheric functional connectivity in rhesus monkeys. Proceedings of the National Academy of Sciences. 2013;110(34):13982–7.

16. Turchi J, Chang C, Frank QY, Russ BE, David KY, Cortes CR, et al. The basal forebrain regulates global resting-state fMRI fluctuations. Neuron. 2018;97(4):940-52. e4.

17. Wang Y, Zheng Y, Xu X, Dubuisson E, Bao Q, Lu J, et al. Electrochemical delamination of CVD-grown graphene film: toward the recyclable use of copper catalyst. ACS nano. 2011;5(12):9927–33.

18. Thunemann M, Lu Y, Liu X, Kılıç K, Desjardins M, Vandenberghe M, et al. Deep 2-photon imaging and artifact-free optogenetics through transparent graphene microelectrode arrays. Nature Communications. 2018;9(1):2035.

19. Makino H, Ren C, Liu H, Kim AN, Kondapaneni N, Liu X, et al. Transformation of cortex-wide emergent properties during motor learning. Neuron. 2017;94(4):880–90.

20. Paszke A, Gross S, Massa F, Lerer A, Bradbury J, Chanan G, et al., editors. PyTorch: An imperative style, high-performance deep learning library. Advances in Neural Information Processing Systems; 2019.

21. Liu X, Lu Y, Iseri E, Shi Y, Kuzum D. A Compact Closed-Loop Optogenetics System Based on Artifact-Free Transparent Graphene Electrodes. Frontiers in neuroscience. 2018;12:132.

22. Cardin JA, Carlen M, Meletis K, Knoblich U, Zhang F, Deisseroth K, et al. Targeted optogenetic stimulation and recording of neurons in vivo using cell-type-specific expression of Channelrhodopsin-2. Nature Protocols. 2010;5(2):247–54.

23. Lu Y, Liu X, Hattori R, Ren C, Zhang X, Komiyama T, et al. Ultralow impedance graphene microelectrodes with high optical transparency for simultaneous deep two-photon imaging in transgenic mice. Advanced Functional Materials. 2018;28(31):1800002.

24. Wekselblatt JB, Flister ED, Piscopo DM, Niell CM. Large-scale imaging of cortical dynamics during sensory perception and behavior. Journal of neurophysiology. 2016;115(6):2852–66.

25. Ren C, Komiyama T. Characterizing Cortex-Wide Dynamics with Wide-Field Calcium Imaging. J Neurosci. 2021;41(19):4160–8.

26. Liu X, Ren C, Lu Y, Liu Y, Kim J-H, Leutgeb S, et al. Multimodal neural recordings with Neuro-FITM uncover diverse patterns of cortical–hippocampal interactions. Nature Neuroscience. 2021.

27. Ma Y, Shaik MA, Kozberg MG, Kim SH, Portes JP, Timerman D, et al. Resting-state hemodynamics are spatiotemporally coupled to synchronized and symmetric neural activity in excitatory neurons. Proceedings of the National Academy of Sciences. 2016;113(52):E8463–E71.

28. Dubey A, Ray S. Cortical Electrocorticogram (ECoG) is a local signal. J Neurosci. 2019;39(22):4299–311.

29. Lindén H, Tetzlaff T, Potjans TC, Pettersen KH, Grün S, Diesmann M, et al. Modeling the spatial reach of the LFP. Neuron. 2011;72(5):859–72.

30. Łęski S, Lindén H, Tetzlaff T, Pettersen KH, Einevoll GTJPcb. Frequency dependence of signal power and spatial reach of the local field potential. 2013;9(7):e1003137.

31. Nunez PL, Srinivasan R. Electric fields of the brain: the neurophysics of EEG: Oxford University Press, USA; 2006.

32. Buzsaki G, Anastassiou CA, Koch C. The origin of extracellular fields and currents -EEG, ECoG, LFP and spikes. Nat Rev Neurosci. 2012;13(6):407–20.

33. Rogers N, Hermiz J, Ganji M, Kaestner E, Kılıç K, Hossain L, et al. Correlation structure in micro-ECoG recordings is described by spatially coherent components. PLoS computational biology. 2019;15(2):e1006769.

34. Muller L, Hamilton LS, Edwards E, Bouchard KE, Chang EF. Spatial resolution dependence on spectral frequency in human speech cortex electrocorticography. J Neural Eng. 2016;13(5):056013.

