## Supplementary Material for "Decoding of cortex-wide brain activity from local recordings of neural potentials"

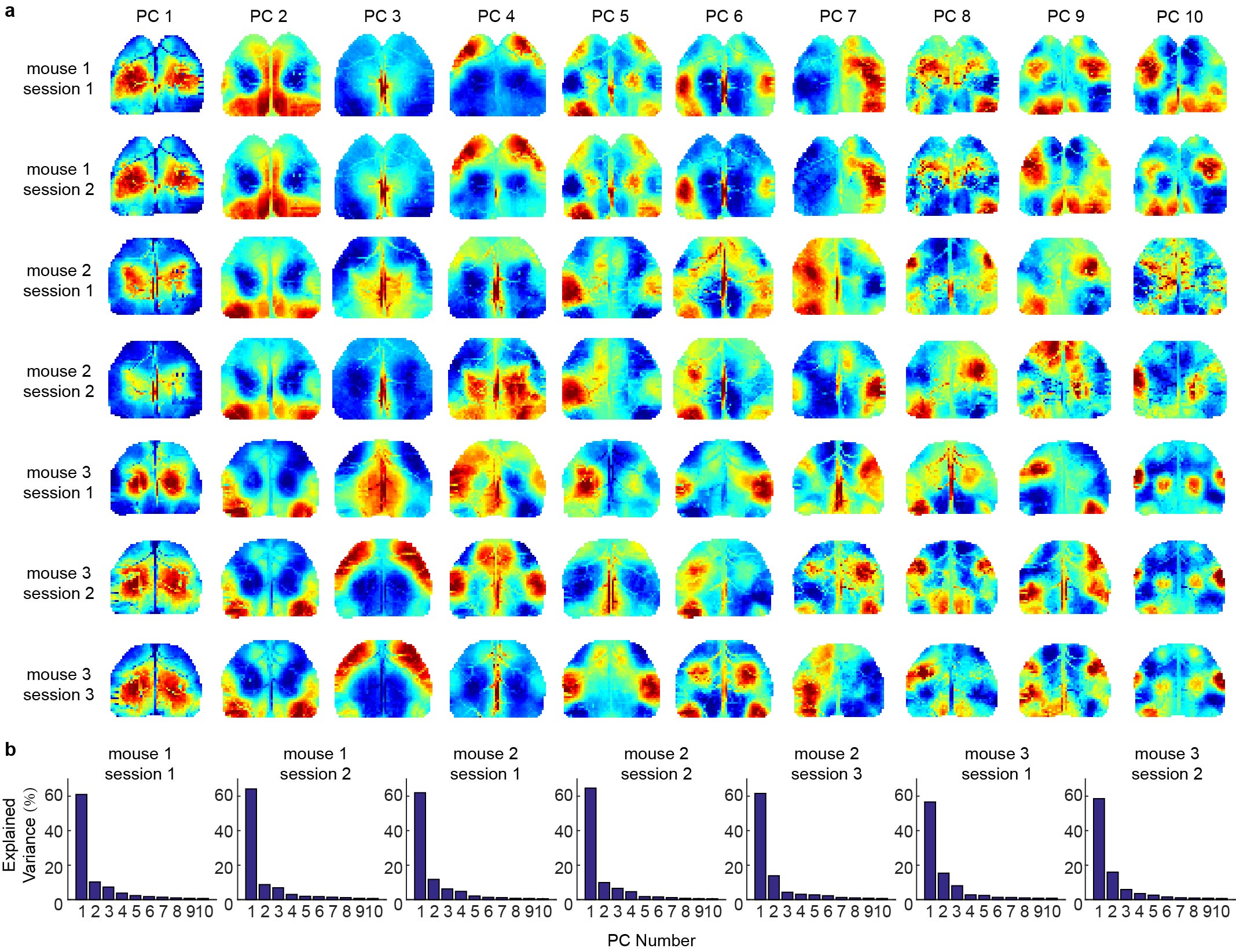

**Supplementary Figure 1. Principal component analysis results for the cortical activity for all the animals.**  (**A)** Identified principal components for individual recording sessions, showing different cortical co-activation patterns. **(B)** The proportion of variance explained by each principal component for individual recording sessions. For all the sessions, the top 10 principal components explained > 92% variance in the data.

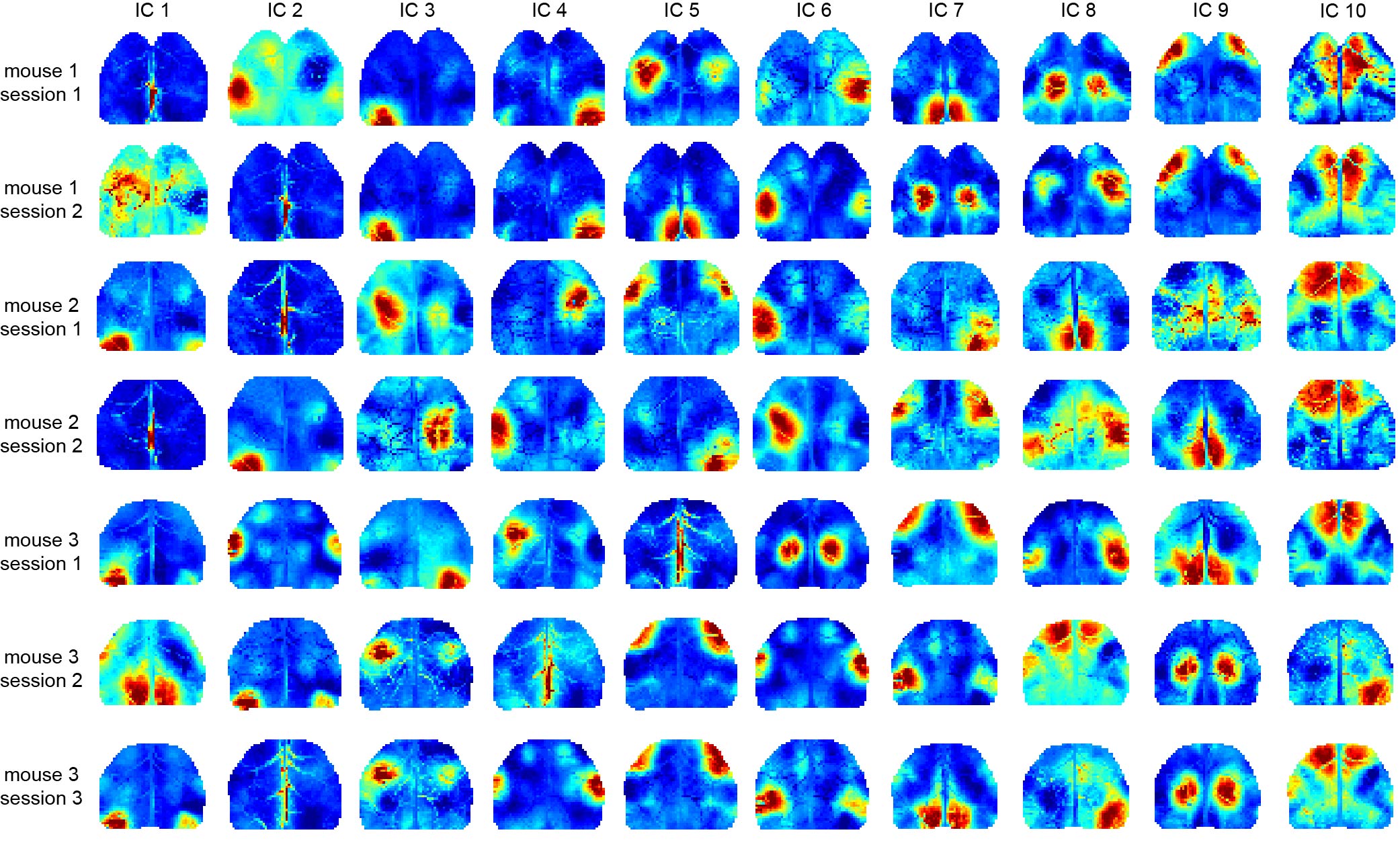

**Supplementary Figure 2. Independent components for cortical activity for all the animals** Similar cortical functional modules and blood vessel activities are identified across different animals.

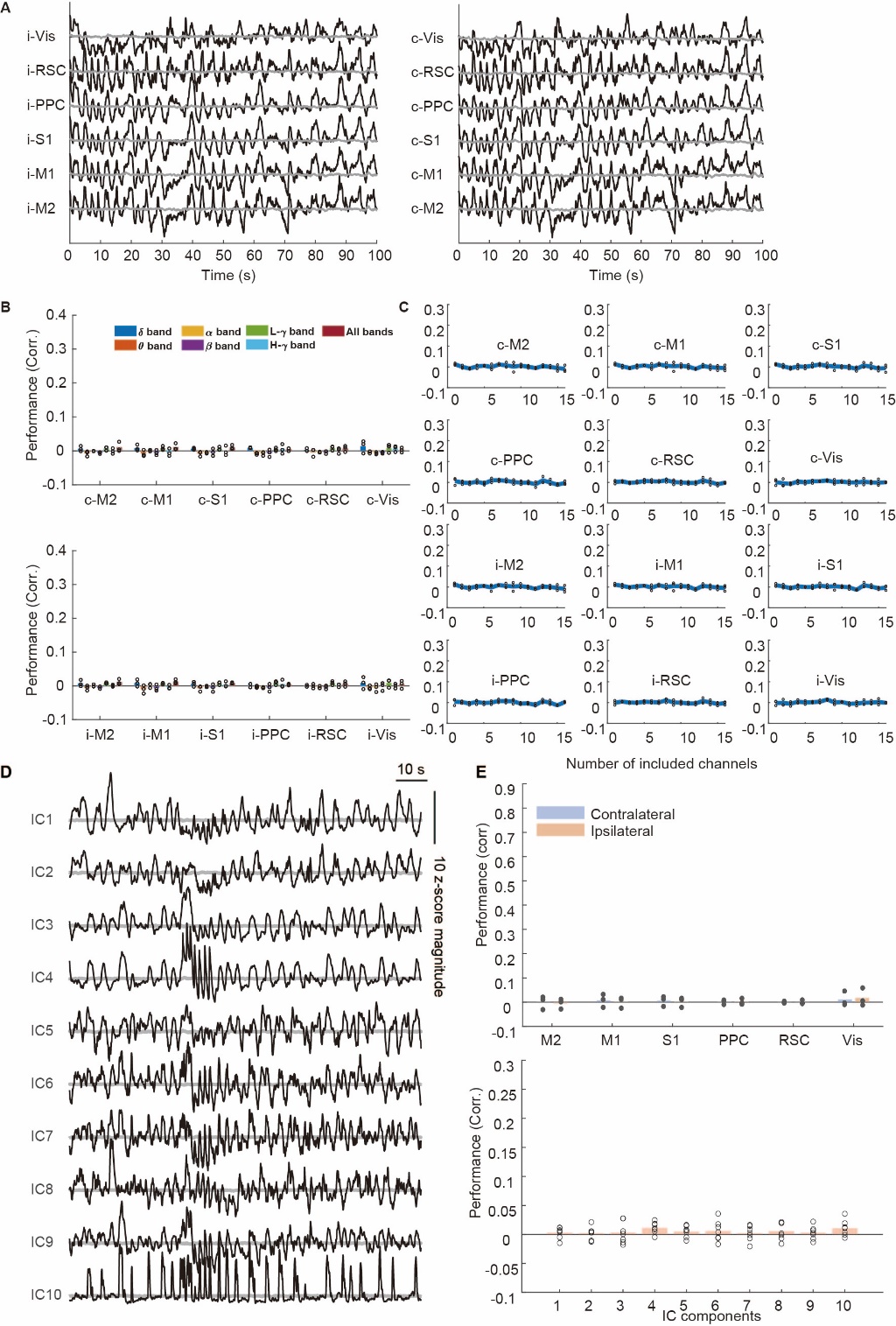

**Supplementary Figure 3. The decoding analysis using shuffled data.** A) The ground truth activity (black) and the decoded activity (gray). B) Decoding performance evaluated for different cortical regions in the contralateral (top) and ipsilateral (bottom) hemispheres using shuffled data from different frequency bands. C) Decoding performance for different cortical regions in the contralateral (top) and ipsilateral (bottom) hemispheres using shuffled data from all the frequency bands, but different numbers of recording channels. D) Decoded (gray) and ground truth (black) weighting scores of the observed cortex-wide activity onto the 10 ICs shown in Figure 4A using shuffled data. E) Decoding performance evaluated at pixel-level for all the cortical regions in the ipsilateral and contralateral hemispheres using shuffled data.

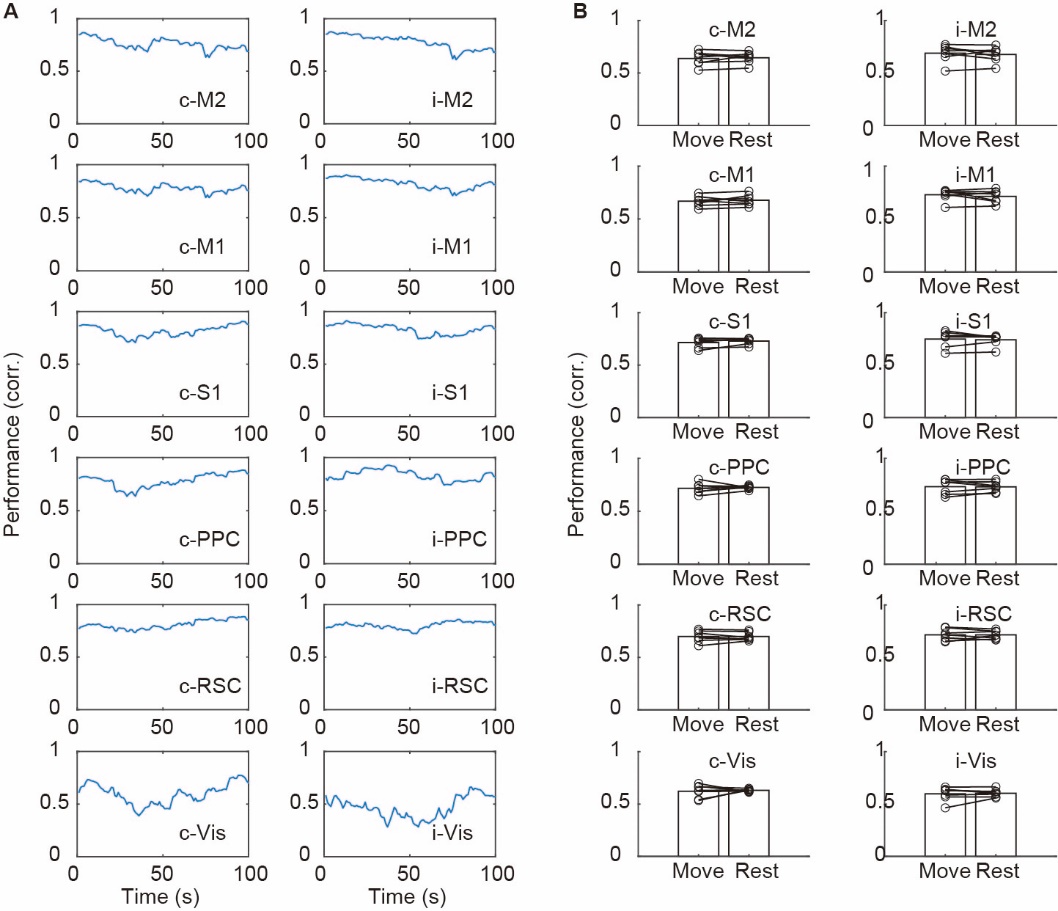

**Supplementary Figure 4. The stability of decoding performance and the effect of movement.** A) The decoding performance evaluated at 30 s sliding window for the data shown in Figure 3A. B) The decoding performance of individual cortical regions between movement and rest phases. Each dot marks the mean correlation evaluated by 10-fold cross-validation using the data recorded from one session.

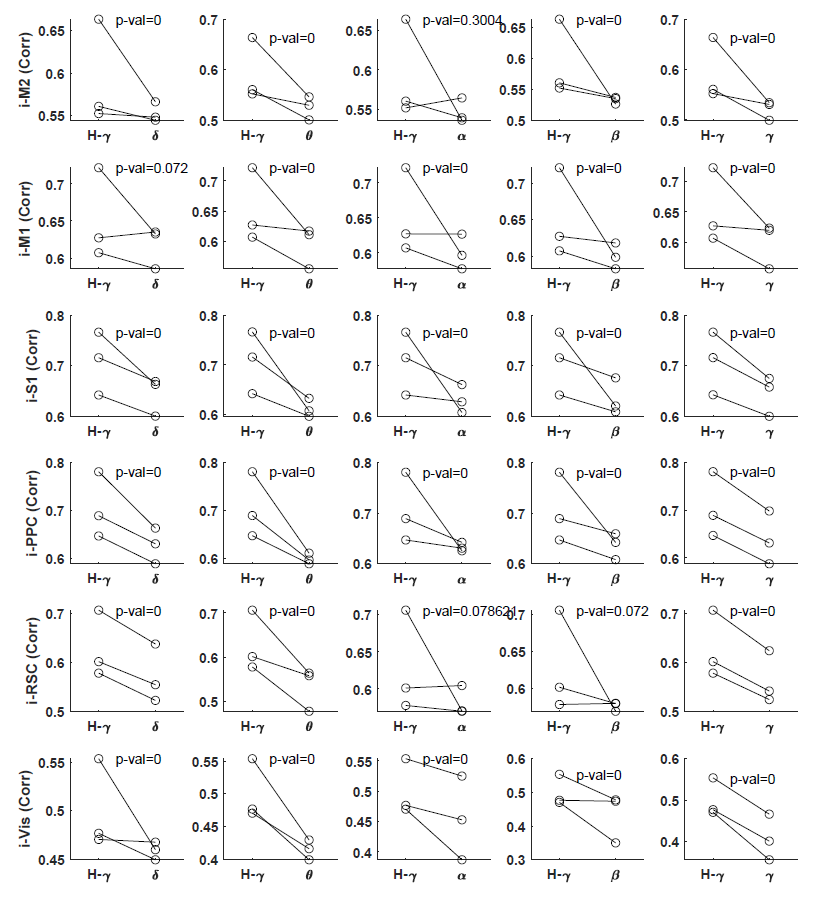

**Supplementary Figure 5.** **Comparison of the decoding performance using High-gamma power band vs. other frequency bands for ipsilateral cortical regions.** For most cortical regions, the high-gamma power band gives significantly higher decoding performance than other frequency bands.

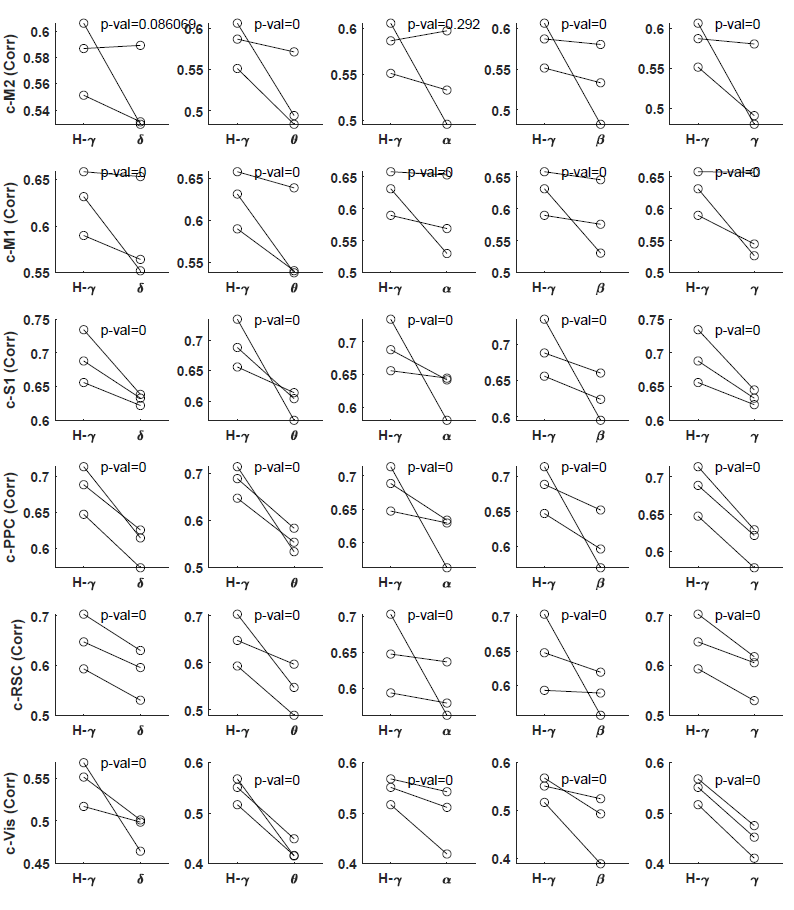

**Supplementary Figure 6**. **Same as Supplementary Figure 5, but for contralateral cortical regions.** For most cortical regions, the high-gamma power band gives significantly higher decoding performance than other frequency bands.

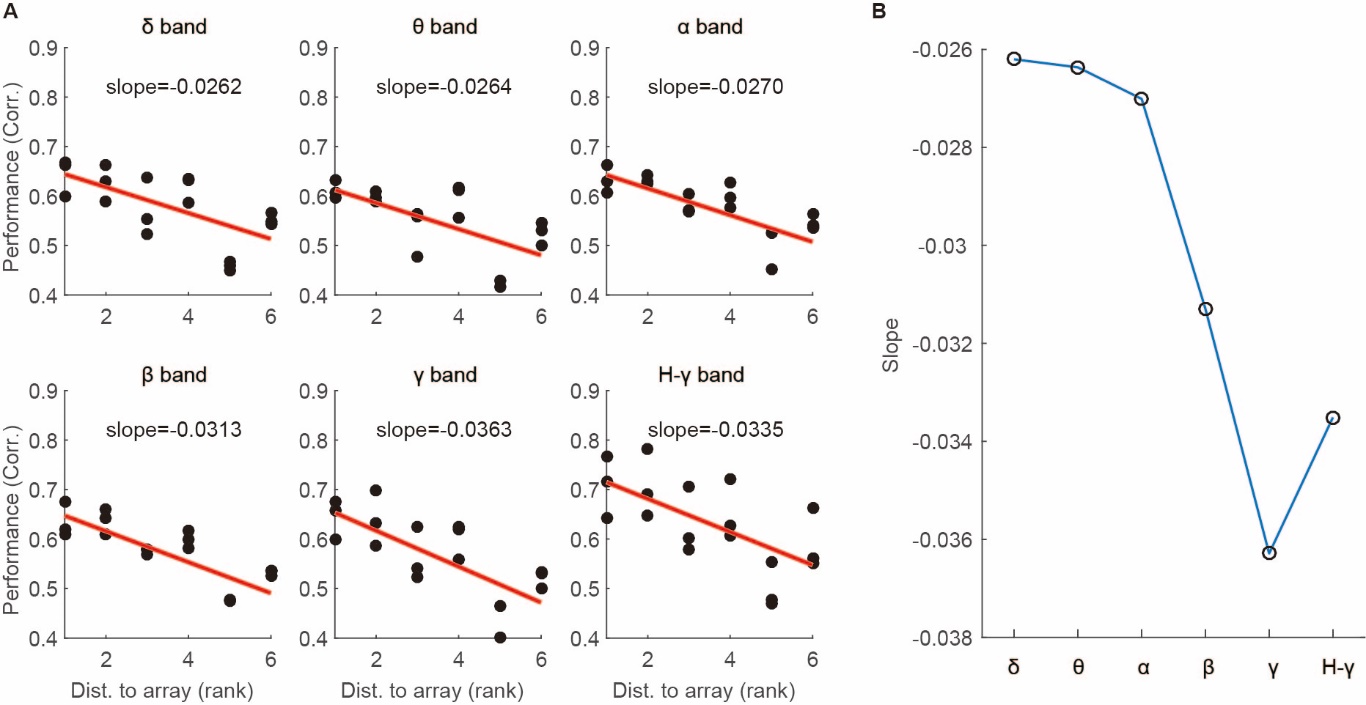

**Supplementary Figure 7. Slope of the decoding performance for different frequency bands.** (A) The decoding performance of ipsilateral cortices plotted against distance rank to the array for different frequency bands. (B) The slope of the decoding performance vs distance for different frequency bands.

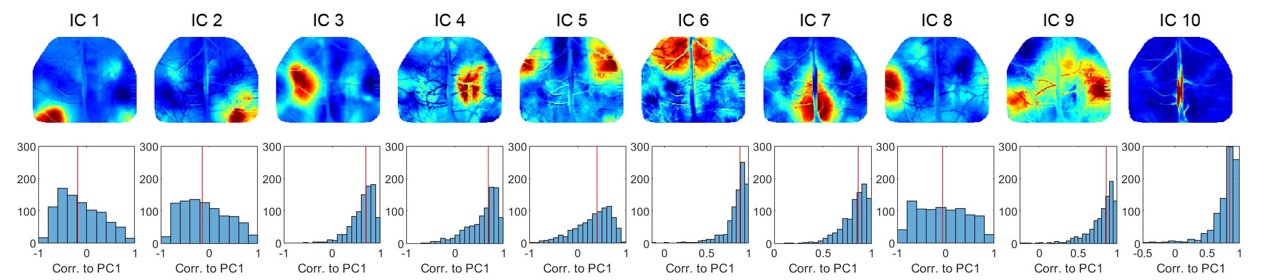

**Supplementary Figure 8. The correlation between the activity of each ICs and the PC1.** Top row shows the template for all the 10 ICs, same as main Figure 4a. Bottom row shows the histogram of the correlation between the activity of each IC and PC1 calculated for nonoverlapping 4s segments during the recording. Note that the IC1, IC2 and IC8 have a median correlation close to zero, showing that their activities are not strongly correlated to PC1.

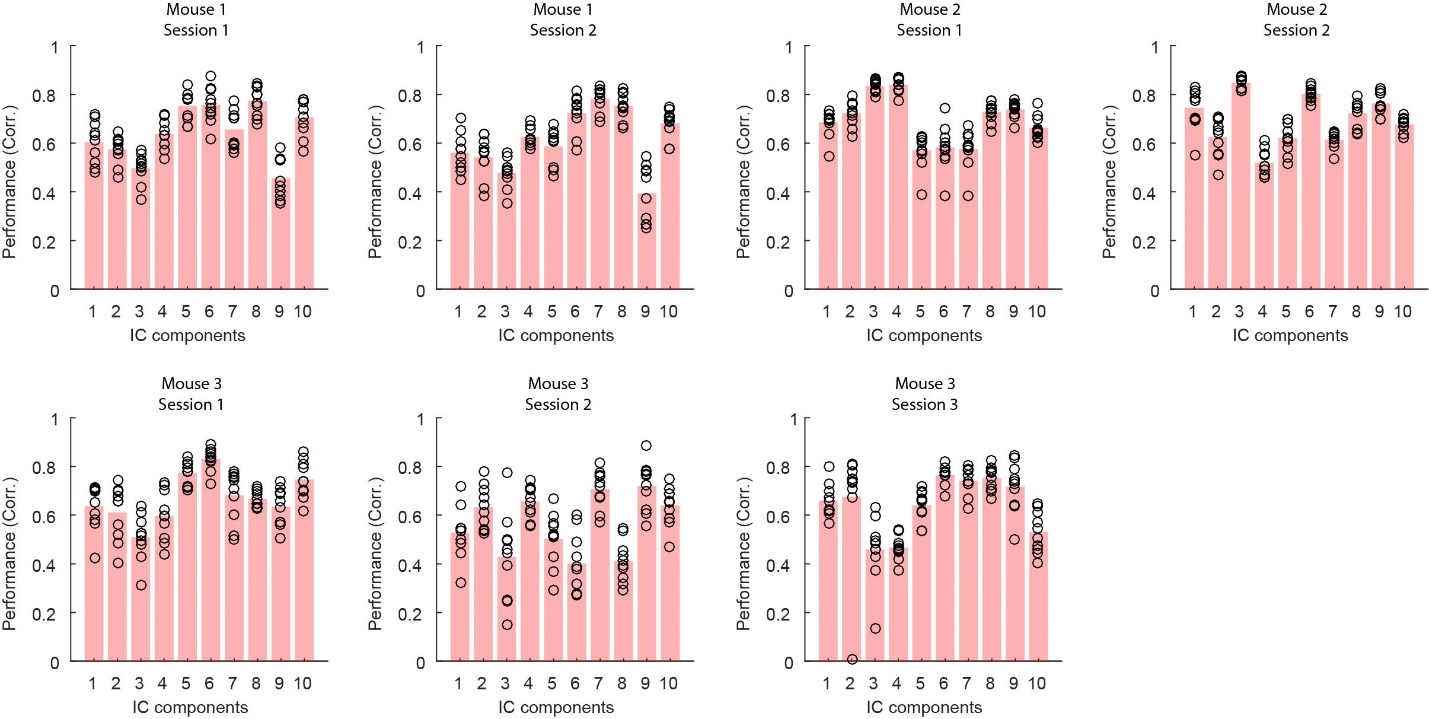

**Supplementary Figure 9. Decoding performance of the IC scores for all the sessions recorded for all the animals.** Each dot marks one cross-validated correlation for one fold data.

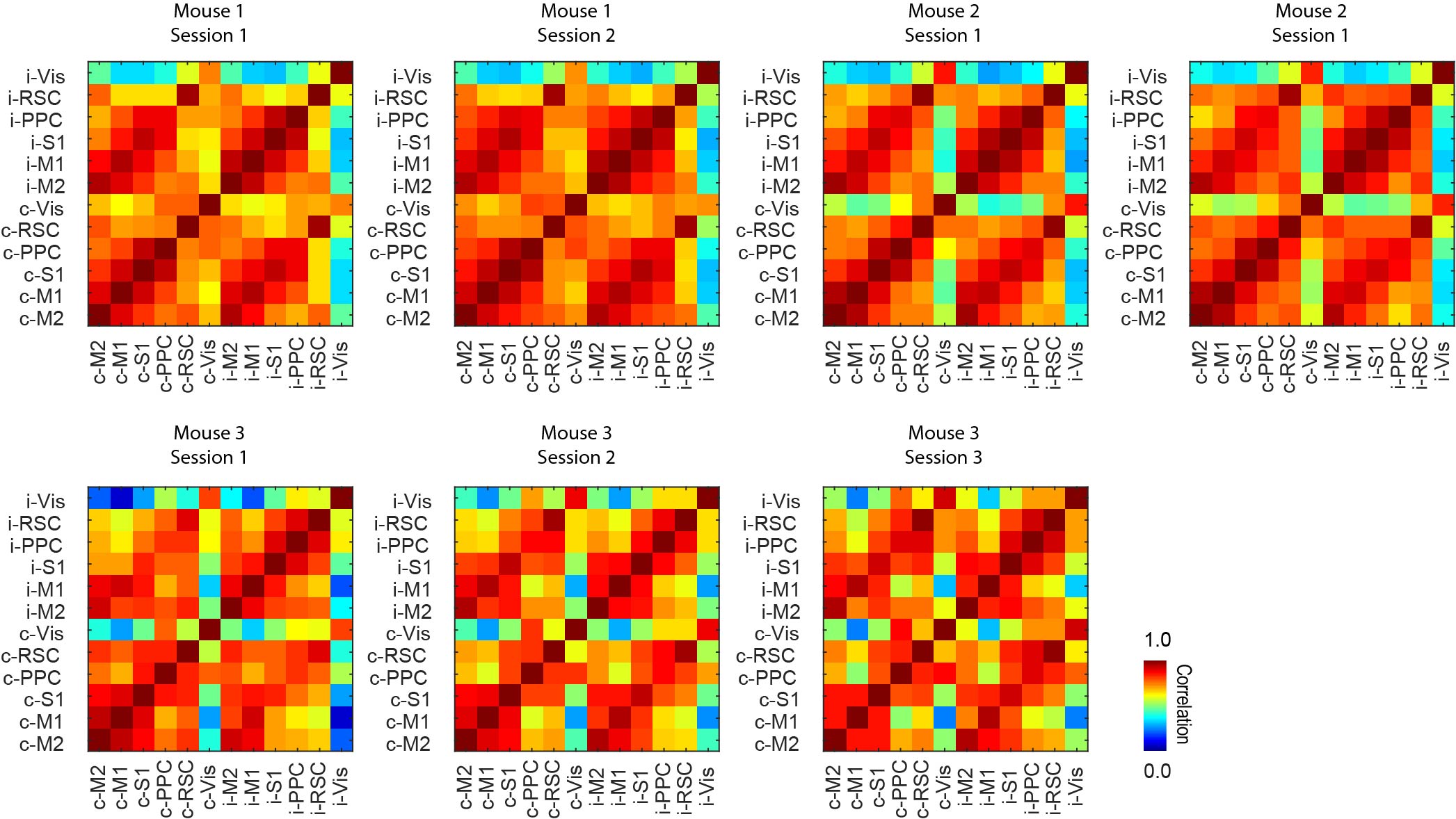

**Supplementary Figure 10. Correlation of ΔF/F activity from different cortical regions in all the mice.** The same functional regions from both hemispheres often exhibit high correlation.

**Movie S1. The decoded (left) and ground truth (right) cortex-wide brain activity evaluated in the held-out data set for Mouse 1 Session 1.** The decoded activities closely resemble the ground truth ones.

**Movie S2. The decoded (left) and ground truth (right) cortex-wide brain activity evaluated in the held-out data set for Mouse 1 Session 2.** The decoded activities closely resemble the ground truth ones.

**Movie S3. The decoded (left) and ground truth (right) cortex-wide brain activity evaluated in the held-out data set for Mouse 2 Session 1.** The decoded activities closely resemble the ground truth ones.

**Movie S4. The decoded (left) and ground truth (right) cortex-wide brain activity evaluated in the held-out data set for Mouse 2 Session 2.** The decoded activities closely resemble the ground truth ones.

**Movie S5. The decoded (left) and ground truth (right) cortex-wide brain activity evaluated in the held-out data set for Mouse 3 Session 1.** The decoded activities closely resemble the ground truth ones.

**Movie S6. The decoded (left) and ground truth (right) cortex-wide brain activity evaluated in the held-out data set for Mouse 3 Session 2.** The decoded activities closely resemble the ground truth ones.

**Movie S7. The decoded (left) and ground truth (right) cortex-wide brain activity evaluated in the held-out data set for Mouse 3 Session 3.** The decoded activities closely resemble the ground truth ones.
