## Supplementary Videos for "Decoding of cortex-wide brain activity from local recordings of neural potentials"

### Slide 1
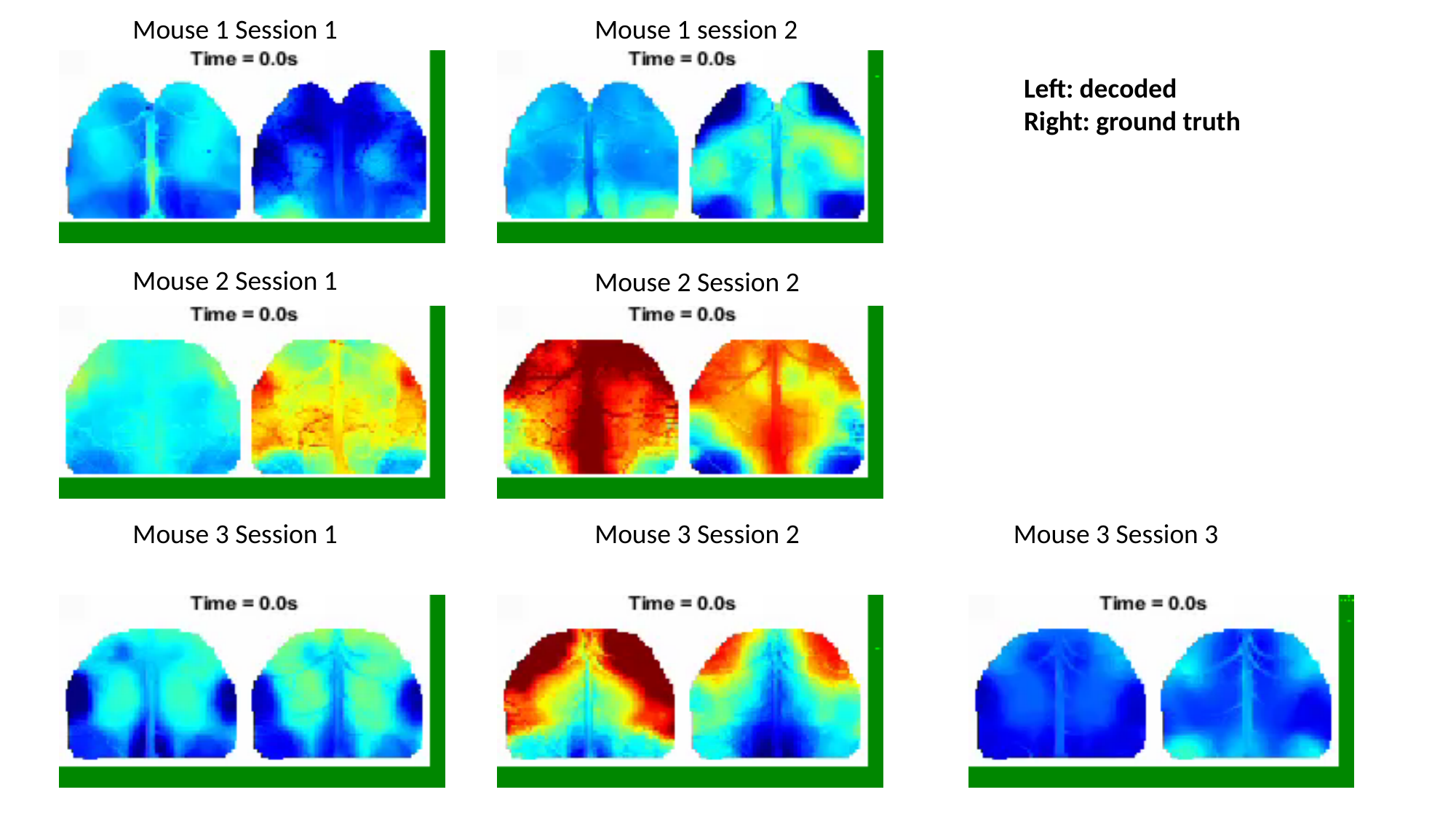

Mouse 1 Session 1
Mouse 1 session 2
Left: decoded
Right: ground truth
Mouse 2 Session 1
Mouse 2 Session 2
Mouse 3 Session 1
Mouse 3 Session 2
Mouse 3 Session 3
